# Within-animal comparison of microdialysis and fiber photometry in amphetamine-exposed mice

**DOI:** 10.1101/2022.10.27.513984

**Authors:** Aske L. Ejdrup, Joel Wellbourne-Wood, Jakob K. Dreyer, Nina Guldhammer, Matthew D. Lycas, Ulrik Gether, Benjamin J. Hall, Gunnar Sørensen

## Abstract

A fundamental concept in neuroscience is the transmission of information between neurons via neurotransmitters, -modulators, and – peptides. For the past decades the gold standard for measuring neurochemicals in awake animals has been microdialysis (MD). The emergence of genetically encoded fluorescence-based biosensors, as well as in vivo optical techniques such as fiber photometry (FP), have introduced technologically distinct means of measuring neurotransmission. To directly compare MD and FP, we performed concurrent within-animal recordings of extracellular dopamine (DA) in the dorsal striatum (DS) before and after administration of amphetamine in awake, freely behaving mice expressing the dopamine sensor dLight1.3b. We show that despite temporal differences, MD- and FP-based readouts of DA correlate well within mice. Down-sampling of FP data showed temporal correlation to MD data, with less variance observed using FP. We also present evidence that DA fluctuations periodically reach low levels and naïve animals have rapid, pre-drug DA dynamics measured with FP that correlate to the subsequent pharmacodynamics of amphetamine as measured with MD and FP.

## Introduction

Monitoring the concentration of extracellular signalling molecules is important to understand brain function and pathology. Several methods with differences in selectivity, resolution, and duration have been used to investigate the characteristics of neuronal signalling. Electrochemical methods such as fast-scan cyclic voltammetry (FSCV) can achieve sub-second temporal resolution, but chemometric data processing restricts the total duration of the measurement to a few minutes for most practical purposes. FCSV offers some molecular specificity but requires the molecule to be electroactive and free from interference from other molecules with similar electrochemical properties (***Jaquins-Gerstl and Michael, 2015***; ***Rodeberg et al., 2017***). In comparison, microdialysis (MD) provides a lower sampling rate in the range of many minutes to hours (***Chefer et al., 2009***), although faster rates have been described (***Wang et al., 2008***). An advantage of MD, however, is its high specificity for molecular species and their metabolites via analysis of dialysates using high performance liquid chromatography (HPLC) or mass spectrometry (MS). Traditionally, MD experiments are performed over hours in total duration. Samples are collected via slow flow through a probe with a diameter in the range of 220-300 μm for mouse probes.

Genetically encoded biosensors offer an optical method for monitoring neurotransmission in-vivo (***Dong et al., 2021***; ***Duffet et al., 2022***; ***Patriarchi et al., 2018***; ***Unger et al., 2020***). The fluorescent signal from these biosensors is typically sampled at a time resolution comparable to or higher than FSCV (***Salinas et al., 2022***), and offers the ability to capture changes in neurotransmitter levels over longer timescales. Biosensors for a host of molecules have been developed, making it a versatile and popular strategy for monitoring in-vivo changes in neuronal signalling (***Sabatini and Tian, 2020***; ***Wu et al., 2022***).

A weakness of data from biosensors, however, is its relative nature. Where MD readily provides measures of concentration, through analysis of dialysates using HPLC or MS, and FCSV can be calibrated ex vivo to provide changes in concentration, biosensors provide relative changes in transmitter concentration. In fact, many FP experiments report changes in z-scored fluorescence levels relative to a fluctuating baseline (***Bian et al., 2022***; ***Robinson et al., 2019***). While this enables easy and reproducible analysis, biological information is lost in z-scoring and individual differences in baseline might be influencing the interpretation of subsequent sections of data.

The aim of this report was to directly compare extended FP recordings of the dopamine (DA) biosensor dLight1.3b fluorescence to DA measurements derived via MD and analysed using HPLC. We sought to use the selectivity and calibrated nature of MD recordings to enhance our understanding of DA neurotransmission as observed through FP. Further, we investigated if this information could provide new insights into DA neurotransmission in response to administration of amphetamine. Our results from the dorsal striatum of both hemispheres of single mice showed that both the time course and peak of amphetamine-induced DA release correlate well between MD and area under curve (AUC) of the FP time series. Notably, FP data showed lower within-group variance. We found that peak DA levels were primarily driven by increases in release rate, although decay of transients in the FP signal was affected as well. Using the correlation established between the two methods from the amphetamine administration, we infer that striatal DA at baseline frequently reached levels below the detection-limit of our FP setup, if only for sub-second intervals. Lastly, our analysis of the FP signal showed that peak DA release rates, pre-injection, correlate with the magnitude of response induced by amphetamine as measured by both MD and FP, highlighting the power of insights into rapid temporal dynamics of extracellular neurotransmitter concentrations.

## Results

### Concurrent MD and FP recordings

To directly compare measurements of extracellular DA between MD and FP we performed concurrent recordings in the dorsal striatum of opposing hemispheres in the same mice (Figure 1A-C). We achieved this by implanting a MD guide cannula containing a dummy probe in one hemisphere. In the opposite hemisphere of the same animal, we injected an adeno-associated virus (AAV) encoding dLight1.3b (***Patriarchi et al., 2018***) expressed under control of the human synapsin promoter (AAV9-hSyn-dLight1.3b) along with implantation of a 200 μm optical fiber for fluorescent signal recording followed by three to four weeks of recovery. On the day of the experiment, we exchanged the dummy probe for an MD probe protruding an additional 1 mm from the guide cannula into the tissue and attached the optical fiber to the FP lens. Mice were habituated for three hours in a circular open arena of 14.5 cm in diameter before beginning recording. We sampled concurrently with both techniques for an 80-minute baseline period, then injected the mice with 1.5 mg/kg amphetamine subcutaneously (s.c.). We then placed them back in the arena for 120 minutes (Figure 1D). Data analysis was limited to 60 minutes prior to injection due to initial photobleaching. Amphetamine was chosen as a circuit activator because of its well-documented effect on extracellular DA in dorsal striatum of rodents and due to its historical role as a pre-clinical model of psychosis (***Butcher et al., 1988***; ***Steinkellner et al., 2014***). Following the in-life phase of the experiments, the MD samples were analyzed on HPLC with a lower limit of detection of 0.25 nM. In vitro recovery of DA across the MD probe membrane was determined in a satellite group to be 8.3 % ± 1.1. From this we inferred a baseline value of 12.9 nM ± 1.8 for the saline group and 10.5 nM ± 1.4 for the amphetamine group for a combined 11.8 nM ± 1.2 across the cohorts. These values are similar to those from previous literature reports (***Sulzer et al., 2016***). Lastly, fiber photometry signals were preprocessed and converted to z-scored dF/F_0_ (zF) for cross-animal comparison (Figure S1A-C and methods).

**Figure 1.**
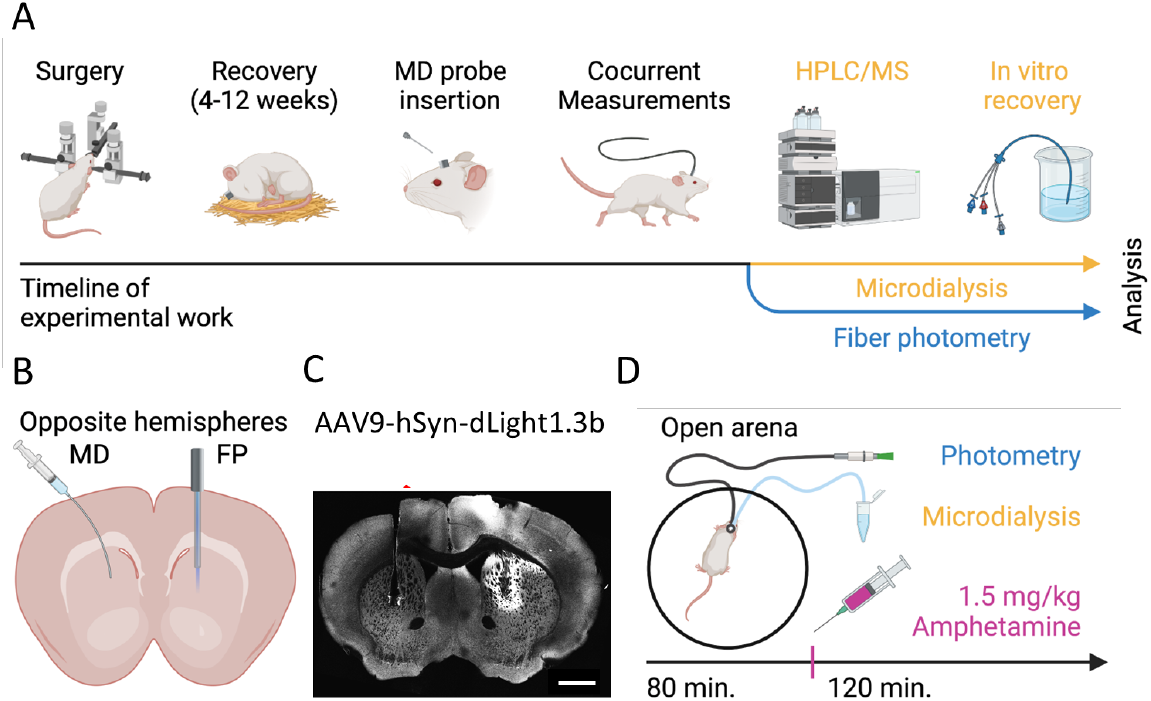
Concurrent within-mouse microdialysis and fiber photometry recordings (A) Schematic for experimental setup. Mice undergo surgery for both MD and FP measurements. Three hours before concurrent recordings an MD probe is inserted. MD dialysate is analyzed using HPLC and baseline values are corrected following in vitro recovery. FP data can be analyzed immediately after acquisition. (B) Sketch of recording location paradigm. Hemisphere placement of MD cannulas and FP lenses are switched between mice. (C) Representative immunohistochemical image of a coronal brain slice of the mouse striatum showing guide canula and probe placement, left, and endogenous cpGFP fluorescence from dLight 1.3b and lens placement, right. (D) Schematic for the measurement sessions. Mice are placed in a circular open arena for three hours of habituation after which 80 minutes of baseline data is sampled prior to injection of 1.5 mg/kg amphetamine. Following injections another 120 minutes is sampled. MD and FP are sampled simultaneously throughout the experiment.

### FP shows earlier peak DA and low variation across animals in response to amphetamine challenge

We first investigated the overall changes in extracellular DA in response to amphetamine as observed with MD. Figure 2A shows DA levels relative to baseline, with samples offset for visibility. The empty circles indicate samples that did not meet technical thresholds for confident peak detection during HPLC and were not included in the remaining analyses. We found that amphetamine administration led to an increase in concentration of DA in the dialysate of all animals. The variation between animals was large, however, with a two-fold increase of DA relative to baseline at the lowest (Figure 2A, purple), and a nearly 60-fold increase at the highest (Figure 2A, gray). DA in individual mice peaked in dialysate collected either between minute 0 to 20 or minute 20 to 40 after amphetamine injection, with a peak average concentration of 8.07 ± 3.03-fold relative to baseline at minute 0-20 (Figure 2B). When converted to absolute concentrations, amphetamine-exposed animals reached 95.3 nM ± 35.8 (Figure S1D). Overall, our MD results are in accordance with similar experiments reported in literature (***Barbier et al., 2007***; ***Becker, 1990***; ***Song et al., 2019***; ***Steinkellner et al., 2014***).

**Figure 2.**
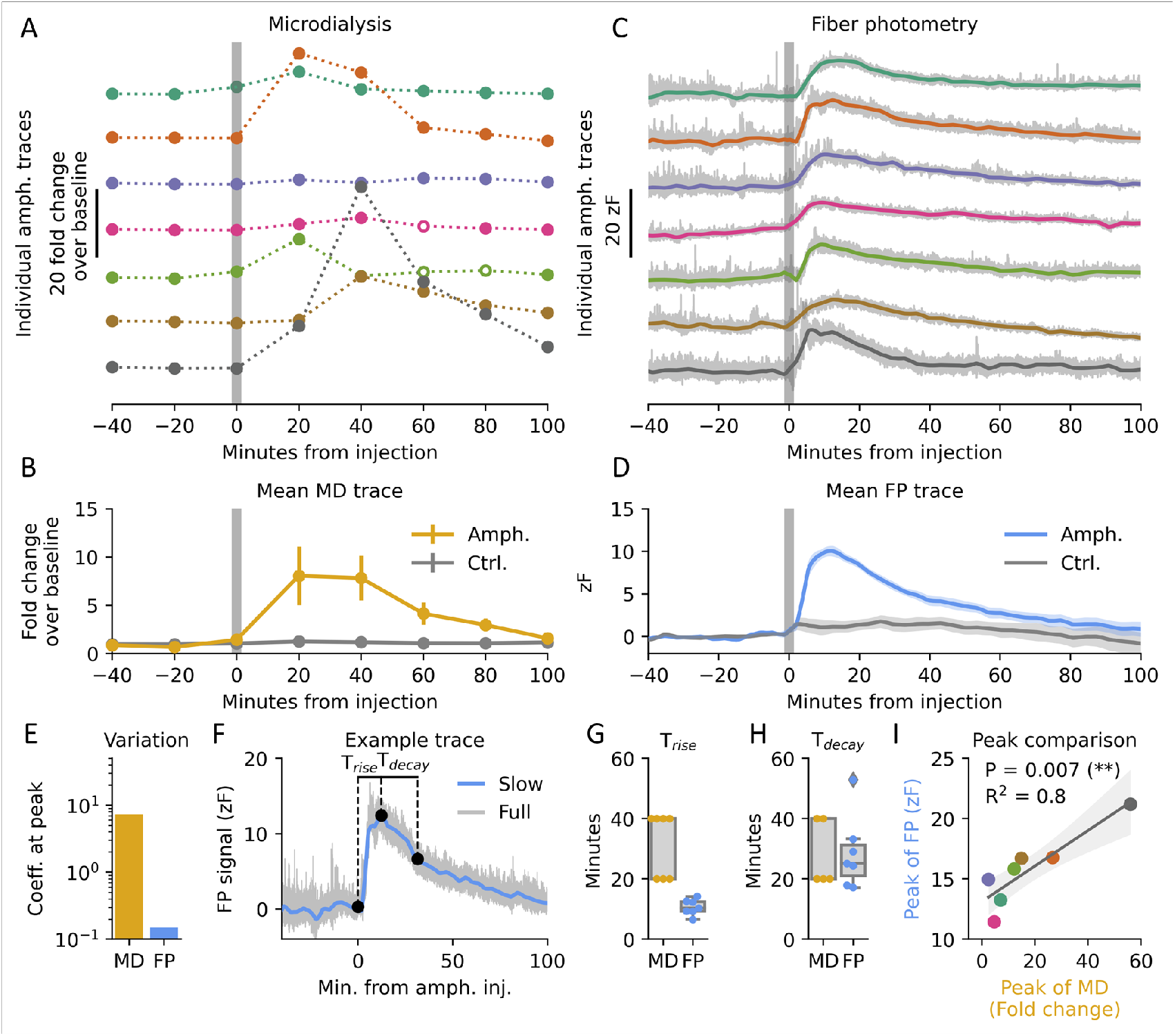
Comparison of amphetamine response (A) MD traces presented as fold change from baseline for all seven amphetamine-injected mice. Hollow points indicate samples that did not pass the signal-to-noise ratio threshold during HPLC analysis of dialysates. (B) Mean MD trace for amphetamine and saline-injected mice. Error bars indicate SEM. (C) Individual FP traces representing dLight1.3b fluorescence for amphetamine-injected mice plotted as z-scored dF/F0 (zF). Thicker lines represent a 0.02 Hz lowpass-filtered signal, with grey background traces showing the unfiltered signal. Colours matching to (A) indicates concurrent recording from same mouse. (D) Mean trace of dLight1.3b fluorescence low-pass filtered at 0.02 Hz for amphetamine and saline-injected mice. Shaded areas indicate SEM. (E) Coefficient variation at peak response to amphetamine for MD and FP. (F) Representative FP trace of dLight1.3b fluorescence lowpass filtered at 0.02 Hz showing quantification of rise time (T_rise_) and decay (T_decay_) of amphetamine response. Unfiltered trace shown as grey background. (G) Rise time of amphetamine response as measured by MD and FP. (H) Half-life of amphetamine response as measured by MD and FP. (I) Peak amphetamine response in MD response correlated to FP dLight1.3b fluorescence. Shaded area indicates 95% C.I. One-sided linear regression, P = 0.003, R^2^ = 0.8, n = 7. Colours match to traces on Figure 2A, C.

We then investigated the concurrent fluorescence response as measured by FP. Figure 2C shows offset response z-scored to pre-injection baseline. We again found a consistent increase in fluorescence after amphetamine, as FP showed a response of 12 to 21 standard deviations above normalized baseline across all animals. Peak fluorescence occurred from 7 to 14 minutes after administration, and the average fluorescence trace peaked after 11 minutes at 10.05 ± 0.56 standard deviations from baseline (Figure 2D). When quantifying variation at peak, we found FP had a coefficient of variation one and a half orders of magnitude lower than MD (Figure 2E). Furthermore, the power spectral density of the FP signal changed markedly after amphetamine, where slower oscillations dominated the signal (Figure S1E-H).

The sub-second temporal resolution of FP allowed us to investigate the kinetics of the response to amphetamine in finer detail than for MD. We defined rise time (T_rise_) as the time from administration to peak of the low pass filtered signal, and decay time (T_decay_) as the time from peak to e-1 (approx. 37%) of peak fluorescence (Figure 2F). These values are poorly determined in MD without advanced interpolative modelling due to the coarse sampling rate. In contrast, they are easily quantified with high precision in FP. For MD T_rise_ was between 0-20 and 20-40 minutes, whereas FP showed a consistent 10.6 ± 1.0 minutes (Figure 2G). Similarly, MD showed a T_decay_ between 20 and 40 minutes, whereas FP had a larger variation than its rise-time, with a decay at 28.5 ± 4.6 minutes (Figure 2H). We observed no correlation between T_rise_ and T_decay_ for FP (Figure S1I). Finally, despite significant methodological differences, we observed a strong correlation of peak amphetamine response between MD and FP, suggesting shared biological information (Figure 2I).

Taken together, we demonstrate a methodology difference when looking at the variance in DA data at peak AMPH effect and T_rise_. No difference between MD and FP was observed when analyzing T_decay_ or when correlating peak MD and FP effect size following AMPH challenge.

### High translatability between MD and FP amphetamine response

So far, we have compared MD and FP with little consideration for what underlying dopamine signal they reflect. MD is seemingly used to capture tonic levels of DA (***Rusheen et al., 2020***), but few studies clearly define what tonic DA is (***Berke, 2018***). Given MD is the most established technique and frequently used to investigate disease states, drugs of abuse, and pharmaceutical interventions (***Darvesh et al., 2011***) we wanted to understand what underlying phenomena are being measured. To that end, we hypothesized that DA measurements obtained from MD reflects area under the curve (A.U.C.) of all DA fluctuations recorded with FP, plus an offset equivalent to the minimum DA levels reached. In other words, MD captures phasic DA as well as any potential minimum basal levels. To approach a direct comparison, we downsampled our FP signal to the same rate as our MD measurement by taking A.U.C. of the preceding 20 minutes as a single datapoint (Figure 3A). While the MD measurements had a much higher spread, in line with findings on Figure 2E, the two methods exhibited a significant linear correlation in response to amphetamine after down-sampling (Figure 3B). Importantly, the two techniques positively correlated in all individual mice (Figure 3C), although not with statistical significance in each (Figure S2A).

**Figure 3.**
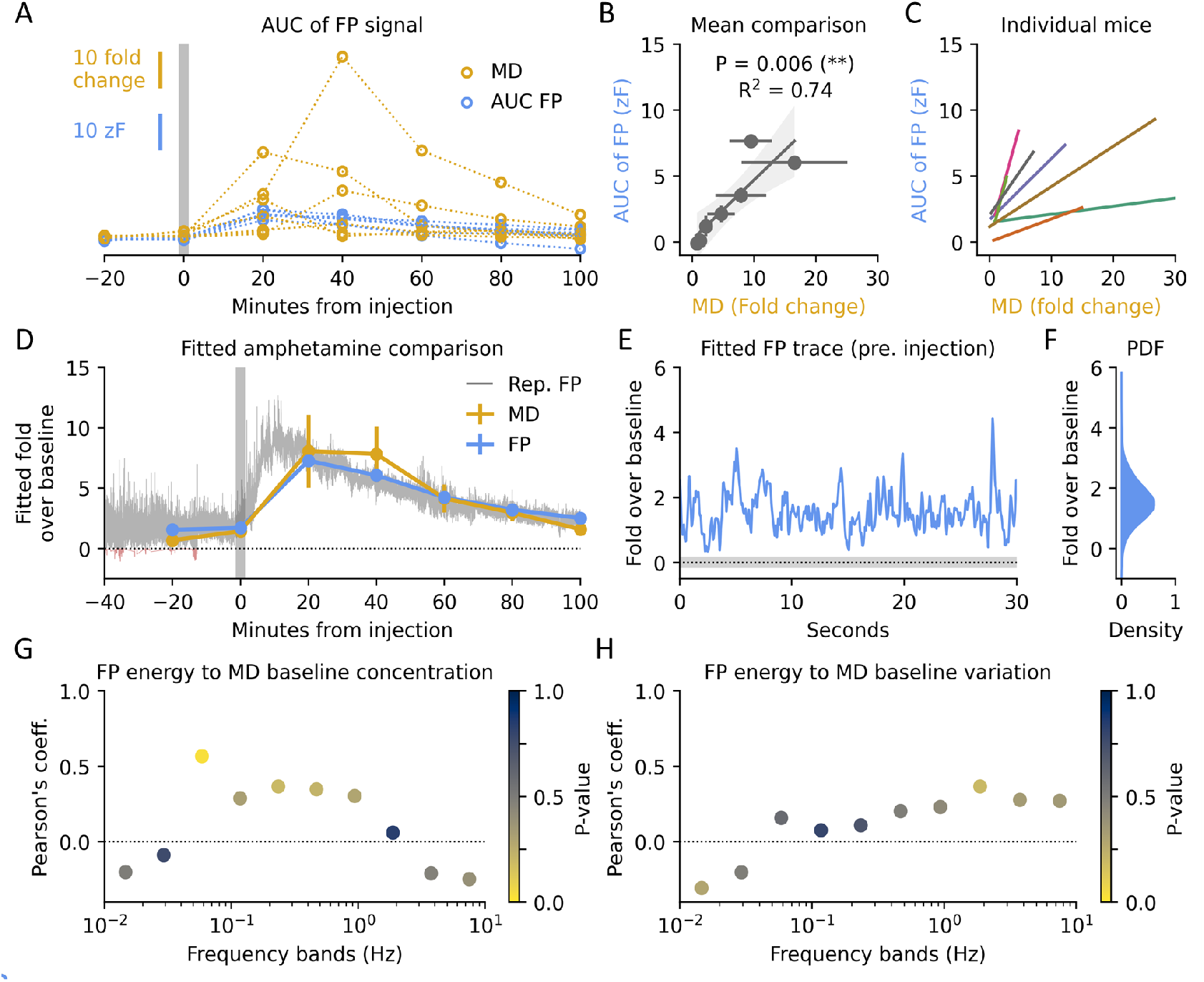
Temporal correlation and baseline estimation (A) Individual MD traces plotted alongside down sampled FP dLight1.3b fluorescence traces by calculating area under the curve of the preceding 20 minutes. (B) Correlation between mean values of MD and down sampled FP. Error bars indicate SEM for individual data points. Shaded area indicates 95% C.I. of the regression. One-sided linear regression, P = 0.006, R2 = 0.74, n = 7. (C) Individual regression lines of transformed FP and MD for each mouse. Colours match traces on Figure 2A, C. Individual statistics and data points can be found on Figure S2A. (D) Mean of down sampled FP during amphetamine injection fitted to MD by linear regression as in (B). Background trace is a representative, fitted, unfiltered FP trace. Error bars indicate SEM. (E) Representative fitted FP trace before injection. Shaded area indicates 95% C.I. of intercept in (B). (F) Probability density function (PDF) of FP values across all mice after application of fit in (B). Only pre-injection data are included. (G) Pearson’s coefficient between MD baseline values and relative energy at different frequency bands in FP traces. Colour indicates statistical significance. No values were significant (α = 0.05) after Bonferroni correction for multiple testing. (H) Pearson’s coefficient between MD baseline variation and relative energy at different frequency bands in FP traces. Colour indicates statistical significance. No values were significant (α = 0.05) after Bonferroni correction for multiple testing.

While FP only records arbitrary fluorescence units, dialysate produced via MD and analyzed using HPLC or MS produce data that can be translated into molar values using a standard curve. Given that the techniques correlate well, we aimed to use information from MD to estimate fold over baseline-values for the FP signal. We used the fit from figure 3B to convert the z-scored FP signal to fold over baseline (Figure 3D). Zooming in on the FP trace pre-injection (Figure 3E), we found the signal fluctuated somewhere between 0 to 4 times above the MD-estimated baseline, often reaching values close to zero. The probability density function of the signal before amphetamine injection is shown in Figure 3F. All in all, FP data presents very similar to MD data when analysed group-wise and at the same time resolution.

Furthermore, in vitro recovery allows for estimating the absolute concentration of the extracellular space surrounding the probe (Fig. S1D). After this scaling the concentration values of the FP signal fluctuated between 0 and 40 nM, with occasional peaks at higher levels (Figure S2B-D). The amphetamine-induced response had a mean concentration peak at 118.8 nM ± 7.2.

### No information about MD baseline in rapid FP signal

MD baseline values (i.e., before injection) vary both across and within subjects. However, fiber photometry only provides a relative measure, not direct information about baseline levels. We hypothesized baseline differences observed in MD might be the result of differential rapid dynamics, such as the frequency or magnitude of DA transients (Figure S3A), occluded by the lower temporal resolution of MD. To probe for shared information between the two methods, we correlated baseline concentration across mice with the relative energy across frequency bands of the FP signal (Figure 3G). No individual band survived multiple testing, but we observed a trend for fluctuations in the FP signal matching conventional DA transients (0.05 to 1 Hz). To further test if this domain in unison significantly correlated with baseline levels, we pooled the energy of the frequency bands but found no statistical significance between the two (Figure S3B). Next, we wanted to gauge if differential fluctuations across frequencies could explain the observed intra-mouse variation in MD during baseline (Figure 3H). No significant frequency bands were identified, and the putative transient domain lost any correlation to the baseline when compared to the variation coefficient (Figure S3C).

Based upon these data we concluded the two techniques do not share immediate shared information at baseline, either due to high variance of MD or lack of information in the FP signal.

### Rapid dynamics at baseline correlate with magnitude of amphetamine response

A strength of biosensors is their ability to measure rapid dynamics at sampling rates even beyond conventional FSCV (***Salinas et al., 2022***). Several studies have shown how these fluctuations encode behavior (***Hamid et al., 2021***; ***Liu et al., 2022***). In our case we wanted to investigate whether rapid dynamics at baseline could have predictive power for future drug response. Mimicking Figure 3G and H, we correlated relative energy across frequency bands of the FP signal to peak MD amphetamine response (Figure 4A). We found statistical significance of a negative correlation in several of the slower domains (< 1 Hz), whereas the most rapid domains correlated positively. As amphetamine is believed to both block DAT recycling of extracellular DA and induce DA release from dopaminergic terminals (***Reith and Gnegy, 2020***), we hypothesized these correlations should be further elucidated by parameters of uptake and release. To that end, we construed two parameter-less measures: the time constant (τ) of the decay in the autocorrelation as an indicator of DAT capacity (Figure 4B), and the 99.5th percentile of the derivate as an indicator of release rate (Figure 4C). We intentionally stayed away from peak identification-based measures, as they require several user-defined input parameters.

**Figure 4.**
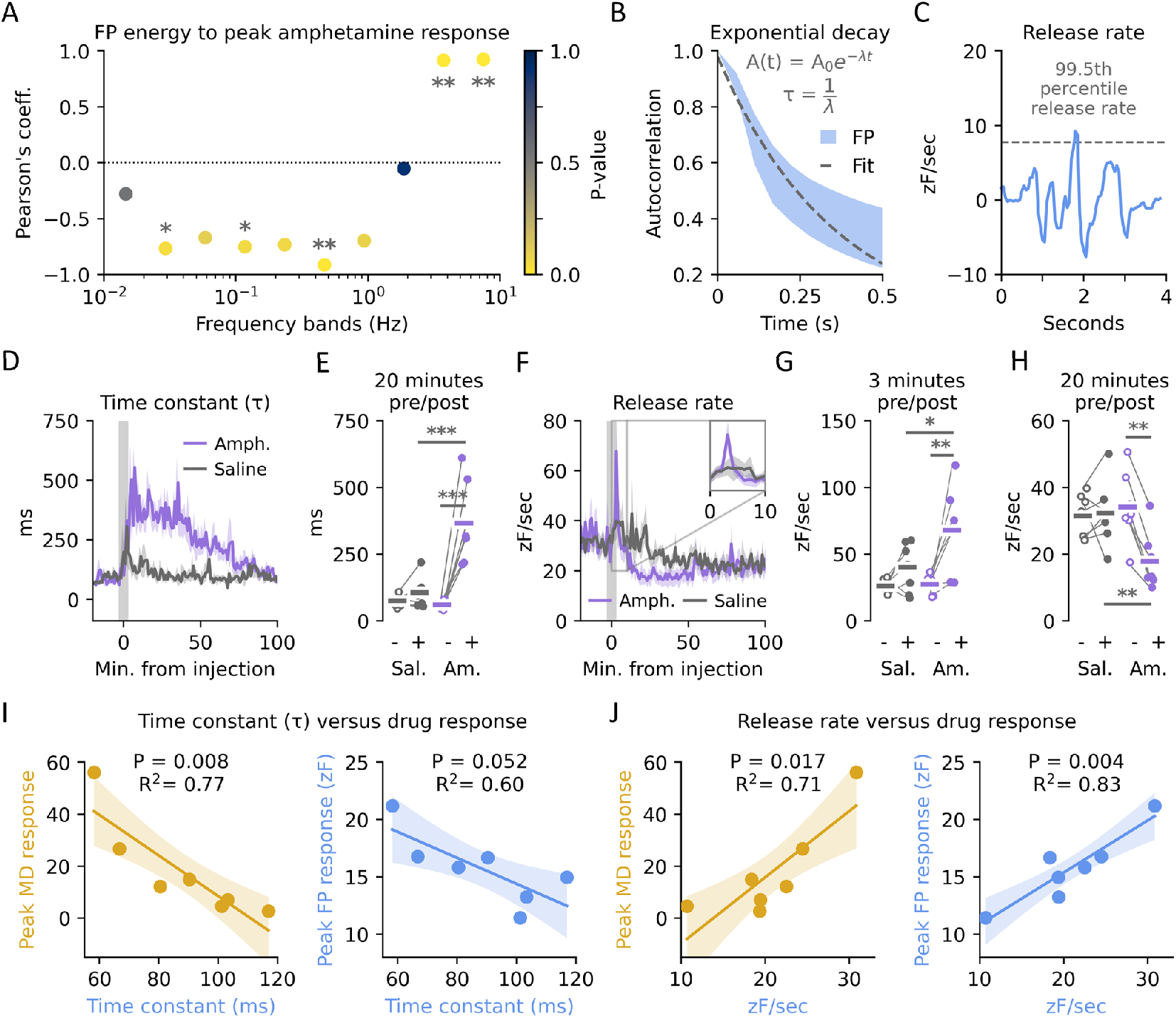
Rapid dopamine dynamics and correlation to drug effects (A) Pearson’s coefficient between peak amphetamine response in MD and relative energy at different frequency bands in FP traces. Colour indicates statistical significance. Two-sided t-test corrected for multiple testing with Bonferroni, n = 7 per test, (*) p < 0.05, (**) p < 0.01. (B) Schematic of the quantification of the time constant (τ). An exponential decay is fitted to the autocorrelation and τ isolated from the inverse of λ. Shaded area indicate SEM of autocorrelation for all traces pre injection and dashed line a representative exponential decay fit. (C) Representative first order derivative trace. Release rate is set to the 99.5th percentile. (D) Time constant (τ) in one-minute bins for amphetamine and saline-injected mice before and after injection. Inset shows zoom of the first 10 minutes. Shaded areas indicate S.E.M. (E) Time constant (τ) 20 minutes before and after injection with either saline (Sal) or amphetamine (Am). One-sided two-way t-test corrected for multiple testing with Bonferroni, n = 7 per group, saline:pre, p = 0.17; pre:pre, p = 1.00; amph:pre, p = 0.001; amph:saline, p = 0.0003. (F) Release rate in one-minute bins for amphetamine and saline-injected mice before and after injection. Inset shows zoom of the first 10 minutes. Shaded areas indicate S.E.M. (G) Release rate 3 minutes before and after injection with either saline or amphetamine. One-sided two-way t-test corrected for multiple testing with Bonferroni, n = 7 per group, saline:pre, p = 0.47; pre:pre, p = 1.00; amph:pre, p = 0.028; amph:saline, p = 0.034. (H) Release rate 20 minutes before and after injection with either saline or amphetamine. Two-sided two-way t-test corrected for multiple testing with Bonferroni, n = 7 per group, saline:pre, p = 0.42; pre:pre, p = 0.28; amph:pre, p = 0.008; amph:saline, p = 0.01. (I) Linear correlation between time constant and peak MD or FP amphetamine response. MD: P = 0.008, R2 = 0.77, n = 7; FP: P = 0.052, R^2^ = 0.60, n = 7. (J) Linear correlation between release rate and peak MD or FP amphetamine response. MD: P = 0.017, R2 = 0.71, n = 7; FP: P = 0.004, R^2^ = 0.83, n = 7.

Consistent with the ability of amphetamine to block reuptake of extracellular DA through DAT, we found a strong effect on τ after s.c. injection (Figure 4D). Prior to administration we found a τ of approximately 100 ms (Figure 4E), consistent with previously reported values from FP measurements in DS (***Salinas et al., 2022***; ***Wei et al., 2022***). Within 5-10 minutes of administration τ rose to 300-400 ms, whereas saline-injected mice only showed a transient response upon injection (Figure 4E). Likewise, 99.5th percentile of the derivate, as a proxy for release rate, was affected by amphetamine (Figure 4F). This effect peaked significantly higher than the saline response after just a few minutes (Figure 4G), but quickly subsided and then undershot the saline injected mice for close to an hour (Figure 4H).

To test if these two parameters could be used in future experiments to predict drug response, we correlated the pre-administration variation between mice to the individual amphetamine response. As the FP traces are z-scored, careful consideration should be exercised – but our MD measurements are free from this caveat. For DAT capacity as a predictor, τ negatively correlated with peak amphetamine response in MD, but fell short of statistical significance when compared to the FP read-out, despite a similar trend (Figure 4I). This negative correlation might be explained by a higher expression of DAT leading to a less efficient inhibition at the sub-saturating concentrations we administered amphetamine at. Conversely, the pre-drug release rate positively correlated with the subsequent amphetamine response (Figure 4J). This correlation was statistically significant for both the MD and FP response, and could reflect a higher propensity for release, either by excess loading of vesicles, release probability, or number of vesicles docked.

In summary, amphetamine affected the rapid DA fluctuations as observed with FP, and both uptake and release parameters of the signal pre-injection correlated with peak amphetamine response as measured by both MD and FP.

## Discussion

Robust measurements of extracellular neurotransmitter concentrations are imperative to improve our understanding of brain function and disease states. These have previously been performed with MD and FSCV, but both methods have limitations with either temporal resolution, neurochemical specificity, or maximal duration of recording. This limits their usefulness in longitudinal within-animal studies necessary to understanding disease progression and treatment. Recently, a new methodological paradigm has been opened by genetically encoded biosensors in combination with invasive optical imaging technologies. When combined, these have potential to capture rapid dynamics over extended periods of time. However, thorough validation and comparison to established methods must be performed to ensure validity of the results.

To contribute to this, we directly compared FP recordings of the dopamine sensor dLight1.3b to MD, as this technique has been the gold standard for assessing neurotransmitter dynamics over time scales of many minutes to hours. We did this by performing concurrent, within-mice measurements of extracellular DA in the dorsal striatum, before and after exposure to amphetamine or saline. While the two measures correlated well on a minutes-to-hours timescale after down-sampling of FP, several parameters of interest could be extracted via the biosensor that were not achievable with MD. Importantly, the rapid dynamics observed have previously been shown to correlate well with FSCV for most applications (***Salinas et al., 2022***). Additionally, FP showed a two-orders of magnitude lower variation. This suggests fewer animals might be needed for a significant effect size if FP is used in place of MD.

As in vitro recovery allows us to assess absolute concentrations in MD, we utilized the strong fit between MD and FP to estimate DA concentrations in the biosensor-based photometry measurements. This put the peaks of spontaneous DA transients somewhere between 30 and 60 nM, which fits well with previous reports from FSCV (***Covey et al., 2014***). Given the sensor was expressed under a pan-neuronal promoter, we assume the FP data reflects mean concentration in the extracellular space below the fiber, and do not rule out the possibility that local concentrations in microdomains may be much higher. However, MD and FP share no immediately apparent information of basal levels before injection, making future estimates for other regions or neurotransmitters purely based on FP difficult.

Lastly, we wanted to investigate if rapid dynamics at baseline could predict drug response. Post-hoc analysis of baseline parameters showed release rate positively correlated with maximal DA response as measured by both MD and FP, whereas transient decay positively correlated with amphetamine response in MD. One reason decay may not significantly correlate with FP amphetamine response is the z-scoring of the signal. Wide transients could lead to a more compressed signal after normalization, which affects the subsequent amphetamine response read-out. For this reason, the MD comparison is key to instill trust in the result.

It has not escaped our notice that such an analysis of baseline DA fluctuations could lead to a pre-selection of animals for studies investigating pharmacokinetic or pharmacodynamic properties of compounds in a drug discovery setting. Furthermore, behavioural and cognitive neuroscience often split animals into high and low performers (***Fernando et al., 2012***; ***Verheij et al., 2008***). Basal dynamics such as this might also help pre-partition experimentally interesting cohorts in the future. Finally, provided future improvement in temporal resolution of neurotransmitter measurements in humans, these analytics might extend into a pre-selection of sub-groups within a disease (***la Fougère et al., 2006***) to design more selective and effective clinical trials.

## Methods & Materials

### Surgery

Male C57Bl/6J mice (3-5 months old, Taconic or Janvier) were used. Mice were housed under a 12-hr light/dark cycle under controlled conditions for regular in-door temperature (21±2°C) and humidity (55±5%) with food and tap water available ad libitum.

Mice were anaesthetised with isoflurane (5% induction, 1.5-2% maintenance in O2/N2O) and placed in a stereotaxic apparatus with mouse adaptor for teeth and ears for head fixation (Kopf 1900). 0.1 mL Marcain (2.5 mg/kg) was injected under the shaven scalp for local analgesia before incision. The exposed skull was dried with a sterile swab. 3 holes were drilled: one for anchor screw, one for the MD guide cannula (Charles River), and one for the 200 μm fiber optic lens (Neurophotometrics). The screw was inserted, andAAV9/2-hSyn1-dLight1.3b was injected into the dorsal striatum using a programmable infusion pump (Drummond Nanoject II, 5nL/sec) (Virus injection co-ordinates: +1.18 mm anterior to bregma (AP), ±1.7 mm laterally (ML), alternating between left and right hemispheres between animals, -2.8 (100 nL), -3.0 (300 nL), and -3.2 mm (100 nL) ventral to dura (DV)). Next, the guide cannula was lowered to the following stereotaxic coordinates: +1.18 mm AP, ±1.7 mm ML, -2.8 DV. Next, dental cement (PHYMEP Superbond) was used to secure the guide cannula in place, making sure not to obstruct the bore hole of the other hemisphere, leaving it clear for implantation. After drying, the FP lens was implanted in the opposite hemisphere, in the same bore hole the virus was injected: +1.18 AP, ±1.7 ML, -3.0 DV. The cement cap was expanded to the other hemisphere, cementing the two implants into a single, very stable implant. The surgery was finalized by placing the skin over the base of the implant, secured using a single stitch. After surgery the mouse was placed in a cage under a heat lamp to wake up and then single housed. The animals received antibiotics (Noromox, 150 mg/mL Amoxicillin s.c.) and analgesics (Norodyl, 0.25 pellet containing 2 mg/pellet Carprofen twice daily) for 5 days following surgery. Following a period of 4-12 weeks for biosensor expression a combined MD and FP study was performed.

### In vivo microdialysis and fiber photometry recordings

On the day of the experiment an MD probe (0.3 mm diameter, 1 mm length, Charles River) was inserted through the guide cannula and a fiber optic cable (Doric Lenses) was attached to the lens using a ceramic mating sleeve (Doric Lenses). The microdialysis probe was perfused with filtered Ringer solution (145 mm NaCl, 3 mM KCl, 1 mM MgCl2, 1.2 mM CaCl2; 1 μl / min) throughout the study. The probe was connected to the microdialysis pump (CMA 400) and a fraction collector (810 microsampler, Univentor). The perfusion buffer consists of artificial CSF and the probe is perfused throughout the experiment. Dialysis set-up allows the animal to move freely during the trial in a bowl containing a layer of bedding. Experimental design consisted of collection of brain dialysate samples in 20 min fractions. Prior to the first sample collection the probes had been perfused for 180 min. A total of 9 fractions were sampled (4 basal fractions and 5 post injection fractions) and dialysate DA content was analysed using HPLC detection. At the same time, a fiber photometry signal was recorded through a fiber optic cable connected to a FP3001 (Neurophotometrics). Mice were injected with saline or amphetamine (1.5 mg/kg) s.c.. After the experiments animals were sacrificed, the brains removed and stored for probe placement verification.

### In vitro microdialysis

Five MD probes (0.3 mm diameter, 1 mm length, Charles River) were lowered into Eppendorf tubes containing a solution of 120 nM DA in aCSF. We collected triplicate samples per probe of 20 minutes each at 1 μL/min flow. Concentration of DA in dialysates was determined by means of HPLC with electrochemical detection and in vitro recovery of DA was calculated.

### HPLC

Concentration of DA in dialysates was determined by means of HPLC with electrochemical detection. Samples were stored refrigerated in a CMA/200 microinjector and separation was performed by reverse phase liquid chromatography (ODS 150 × 3.2 mm column) using mobile phase (150 mM NaH2PO4, 4.8 mM citric acid monohydrate, 3 mM dodecyl sulphate, 50 μM EDTA, 8 mM NaCl, 11.3% methanol and 16.7% acetonitrile, pH 5.6) at a pump flow rate of 0.2 ml/min. Electrochemical detection was accomplished using a coulometric detector and a SenCell (Antec); potential set at E1 = 500 mV (Coulochem III, ESA). Limit of detection was 5 fmol/20μL.

### Fiber photometry preprocessing

Preprocessing of the fluorescence traces was done by applying a 5 second median filter to both the 415 nm and 470 nm channel from the start of recording to 2 minutes before the first injection. A linear fit between fluorescence intensities of the two channels was obtained and applied to the isosbestic 415 nm channel to correct for differential bleaching rates. This fit was applied to the entire 415 nm timeseries, and the two signals were converted to dF/F by subtracting the 415 nm channel from the 470 nm and dividing by the 415 nm. Fluctuations between 10 and 20 Hz were filtered due to lack of resolution, as per the Nyquist criterion, by applying a discrete wavelet transform. Lastly, each trace was z-scored to the mean and standard deviation of the 40 minutes leading up to injection.

### Fiber photometry analysis

Slow fluctuations below 0.01 Hz were extracted by applying the discrete wavelet transform with the “sym4” wavelet using the python PyWavelets v1.2.0 package and for the kinetic analysis peaks were found using the python SciPy v1.5.2 package.

To compare MD and FP we computed area under the curve of FP signal in 20-minute bins leading up each MD sampling and divided by time. A linear fit was obtained between all MD and transformed FP samples across mice and was uniformly applied to all transformed FP traces to convert into fold over baseline.

For the baseline comparison energy was computed by summing the squares of the energy across levels and comparing the energy in each level across mice with either the mean basal levels or variance of the basal levels.

### Statistics

The employed statistical analyses are presented in the legends associated with each figure, and multiple testing was corrected for using the Bonferroni-Holmes method where specified. All n-values are individual mice unless otherwise specified. Statistical analyses were carried out with the open-source python packages SciPy v1.5.2, NumPy v1.18.1, and Seaborn v0.11.0. Boxplots show 25th and 75th percentile, with whiskers indicating data up to 1.5 times the interquartile range. Remaining data are plotted as outliers. No statistical methods were used to predetermine sample sizes. Data analysis was not performed blinded but were automated to a degree where the experimenter had no specific impact on outcome. Sessions with severe fiber tangling were excluded from analysis on a qualitative basis, and MD samples with inconclusive peak detection were excluded.

## Data availability

The data from this study are available from the corresponding author upon reasonable request.

## Acknowledgment

We would like to thank Seven Biosciences for the use of dLight1.3b and for productive discussions of the data. This preprint was created using the LaPreprint template (https://github.com/roaldarbol/lapreprint) by Mikkel Roald-Arbøl.

## Author contributions

JW and NG performed the experiments. ALE developed the analyses with help from JKD and MDL. ALE produced the figures. ALE, JW, JKD, BH and GS discussed the data. ALE, JKD, JW and GS wrote the manuscript. ALE, JW, JKD, UG, BH and GS edited the manuscript. UG, BH, and GS supervised the project. All authors critically reviewed the manuscript.

## Supplementary

**Figure S1.**
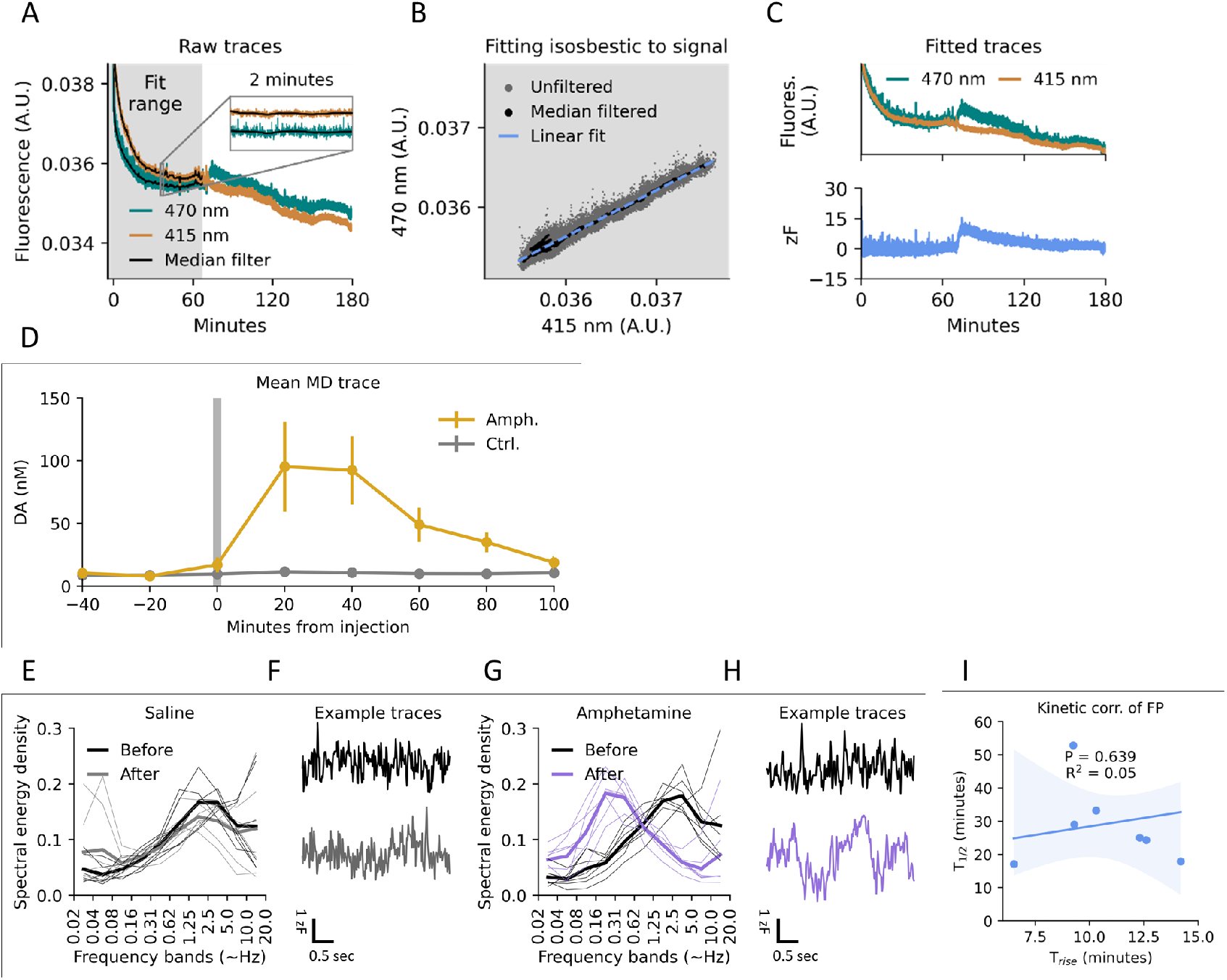
(A) Representative traces of both isosbestic (415 nm) and excitatory (470 nm) channels. Signal is corrected for photobleaching and z-scored from a fit in the grey area. (B) Linear fit between the isosbestic (415 nm) and excitatory (470 nm) channel used to correct for photobleaching. (C) Top: Representative traces after the isosbestic (415 nm) channel has been fitted to the excitatory (470 nm). Bottom: Final signal after subtraction of fitted isosbestic channel (415 nm) and z-scoring. (D) Mean MD traces from Figure 2B converted to DA concentrations after in vitro recovery at 8.3% ± 1.1. (E) Spectral energy density before and after saline injection. Thick lines indicate mean of group. (F) Representative traces of rapid dynamics before and after saline injection. (G) Spectral energy density before and after amphetamine injection. Thick lines indicate mean of group. (H) Representative traces of rapid dynamics before and after amphetamine injection. (I) No significant linear correlation observed between rise time and half-life of amphetamine response measured by FP representing dLight1.3b fluorescence (P = 0.639, R^2^ = 0.047, n = 7). Shaded area indicates 95% C.I.

**Figure S2.**
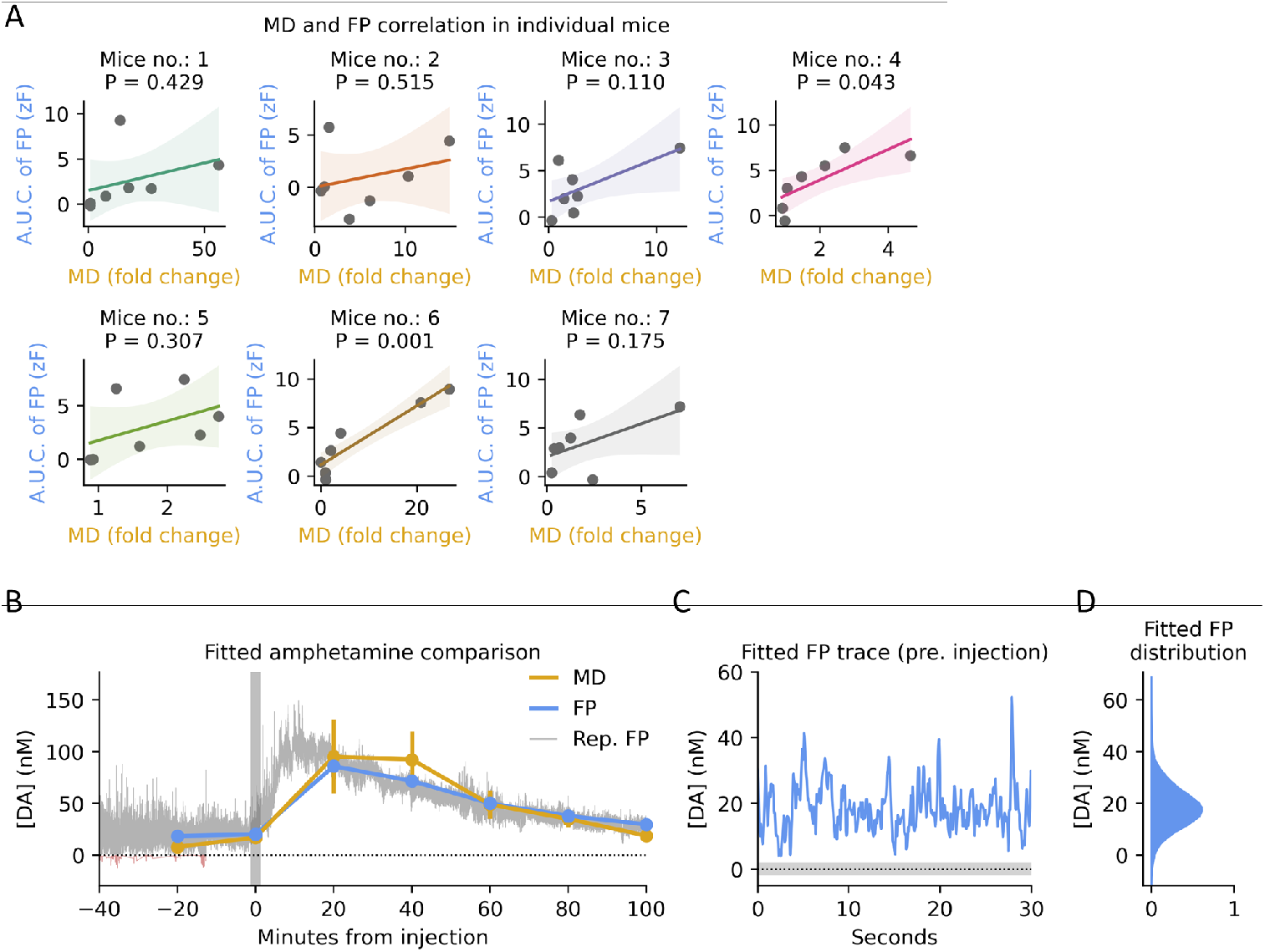
(A) Linear correlation between MD and down sampled FP for each amphetamine-injected mouse. Colours match traces from Figure 2A, C. (B) Mean of down sampled FP during amphetamine injection fitted to MD by linear regression as in Figure 3D converted to DA concentration after in vitro recovery at 8.3% ± 1.1. Background trace is a representative, fitted, unfiltered FP trace. Error bars indicate S.E.M. (C) Representative fitted FP trace before injection as in Figure 3E converted to DA concentration after in vitro recovery. Shaded area indicates 95% C.I. of intercept in (B). (D) Probability density function (PDF) of FP values across all mice after application of fit in Figure 3F converted to DA concentration after in vitro recovery. Only pre-injection data are included.

**Figure S3.**
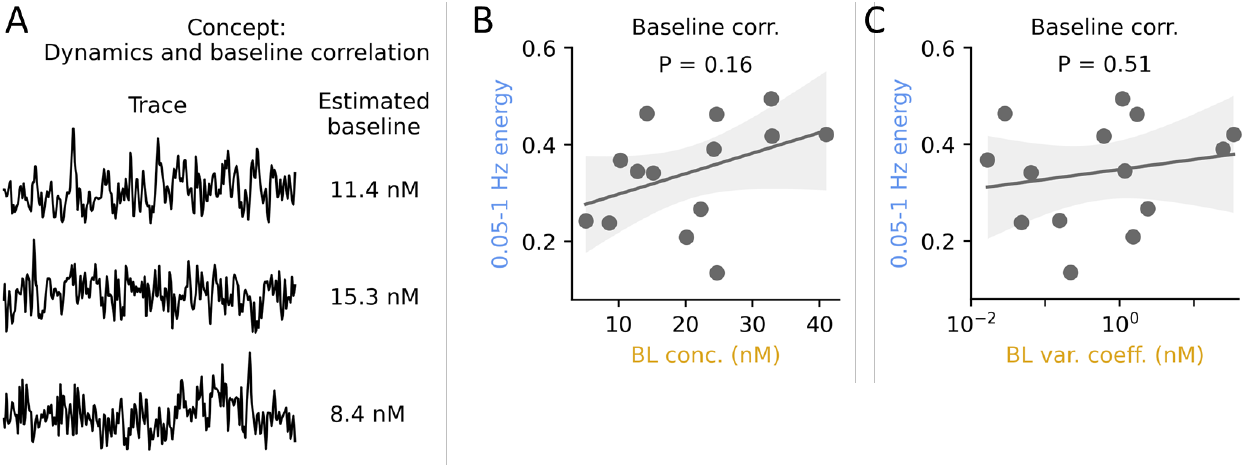
(A) Conceptual schematic of the hypothesis: some attributes in the rapid dynamics of the traces (left) can be used to predict the MD-measured baseline concentration (right). (B) Correlation between MD baseline concentration and energy in putative transient frequency domain. Shaded area indicates 95% C.I. Linear regression, P = 0.16, n = 14. (C) Correlation between MD baseline variation and energy in putative transient frequency domain. Shaded area indicates 95% C.I. Linear regression, P = 0.51, n = 14.

## Notes

### Competing Interest Statement

Joel Wellbourne-Wood, Jakob K. Dreyer, Nina Guldhammer, Benjamin J. Hall and Gunnar Soerensen are all employed at H Lundbeck A/S.

### Summary of Updates

Formatting and reference list updates.

